# Socially integrated female chimpanzees have lower offspring mortality

**DOI:** 10.1101/2025.04.29.651242

**Authors:** Joseph T. Feldblum, Kara K. Walker, Margaret A. Stanton, Elizabeth V. Lonsdorf, Deus Mjungu, Carson M. Murray, Anne E. Pusey

## Abstract

In humans and other social mammals, more socially connected females often have higher fitness. Yet evidence linking female sociality to offspring survival remains inconsistent, and is limited to a handful of primate species in which females depend on close female kin for social status. Here we examine the relationship between female social integration and offspring survival in eastern chimpanzees. We find that females that were more socially integrated with other females in the year before giving birth had higher offspring survival to age 1 (the period of highest mortality) and age 5 (the approximate age of weaning). Furthermore, social integration remained a strong predictor of offspring survival among females without close female kin. Our results thus add to a small set of studies linking sociality with offspring survival, here in the dispersing sex. As in humans, more socially connected female chimpanzees have higher offspring survival, despite primarily residing with non-kin.

## Introduction

In group-living mammals, individuals often form differentiated social bonds and differ in their overall sociality^1^. Recent decades have seen accumulating evidence that sociality is related to health, longevity, and reproductive success in humans and other animals^2–6^. However, among social mammals, evidence for a link between sociality and offspring survival (an important fitness component^7,8^) remains mixed for two reasons. First, evidence for a positive association is taxonomically limited: four studies in cercopithecine monkeys find that more socially integrated females, or those with stronger social bonds, have higher offspring survival (yellow/anubis hybrid baboons, *Papio cynocephalus* with *Papio anubis* admixture^9^; Chacma baboons, *Papio ursinus*^10,11^; vervet monkeys, *Chlorocebus pygerythrus*^12^). But studies in other taxa, as well as recent work in cercopithecines, report no such sociality-offspring survival link (yellow/anubis baboons^13^; chacma baboons^14,15^; bighorn sheep, *Ovis canadensis*^16^; Kinda baboons, *Papio kindae*^17^). In some cases, more social females actually have lower offspring survival (eastern grey kangaroos, *Macropus giganteus*^18^; yellow-bellied marmots, *Marmota flaviventris*^19^), or have higher or lower offspring survival depending on other group dynamics (white-faced capuchins, *Cebus capucinus*^20^).

Second, a recent analysis of data from yellow/anubis hybrid baboons demonstrated evidence of reverse causality in the relationship between female sociality and offspring survival^13^. Females were most social with other females when they had young infants, and with males when cycling, including the cycle immediately following infant death. Controlling for this variation, the authors found no evidence for a positive association between maternal sociality and infant survival^13^. Instead, offspring production and mortality are likely to drive differential female sociality rather than the other way around in this^9,13^ and likely other populations of baboons^10,11,15^. Indeed, in primates and other mammalian species, females often show attraction to, and regularly interact with, other females’ offspring^21–23^, and thus this potential for reverse causation may extend beyond studies of baboons. Of the studies described above, only two^12,13^ use methods that clearly avoid or account for state-dependent social variation as a potential confound, suggesting a critical need for a reevaluation of the relationship between sociality and offspring survival in social mammals.

One reason to expect that female mammals improve their offspring’s survival by being more socially connected is that female sociality could facilitate tolerance or agonistic support, leading to better feeding efficiency or a reduction in received aggression and infanticide in the postpartum period^5,6,24^. Social bonds, tolerance, and agonistic support might be particularly strong among kin due to inclusive fitness effects^25^. On the other hand, because more social females can also incur costs such as feeding competition and increased aggression from males^26,27^, it could be that females in most species may simply not derive sufficient benefit from social bonds to depend on them during the particularly sensitive and energetically costly perinatal period^28,29^. Social organization may also play a role; in species with high fission-fusion dynamics, females have the option to avoid larger foraging parties to reduce the costs of sociality^30^, and could therefore potentially benefit from becoming less social when pregnant or nursing.

Here we investigate the relationship between social integration (here defined as females’ overall affiliative tendency^5^) and offspring survival using 37 years of long-term data on free-ranging eastern chimpanzees (*Pan troglodytes schweinfurthii*) in Gombe National Park, Tanzania. Male philopatry and female dispersal are observed in all known chimpanzee communities, although the degree of female dispersal varies; in Gombe around half of females in one community remain in their natal community to reproduce^31,32^, although only a minority of natal females reside with female kin as adults^33,34^. This dynamic allows us to investigate the role that kinship may play in mediating a link between sociality and offspring survival, and to determine whether there are direct benefits to sociality among females that lack close kin. Chimpanzees also have high fission-fusion dynamics^33^, and in eastern chimpanzees females are less gregarious than are males, more often traveling alone or with only their dependent offspring^35–37^. But female chimpanzees nevertheless form differentiated social relationships with other female^34,38,39^ and sometimes male^40^ community members. Female-female bonds are associated with cooperative behaviors like food sharing^41^ and joint territorial defense^42^, which may impact fitness; the potential adaptive benefits females gain from bonds with males are less clear, but in other primate species these male social partners may provide offspring protection (e.g. ref. ^24^). Females that co-reside with female kin enjoy some fitness-related benefits such as access to higher-quality core areas^33,34^, higher dominance rank^43^, and earlier reproduction^44^. Yet many female bonds are formed with non-kin^34,38^, and females without close kin in their community can still achieve high rank^43^. Therefore this system allows us to examine the influence of dispersal pattern, kin availability, and fission-fusion dynamics on the relationship between sociality and offspring survival.

## Results

### Offspring survival to age 1

We used data from the year before birth (rather than annual or lifetime measures of maternal sociality) to predict offspring survival for two reasons. First, sociality after birth could itself be influenced by offspring presence^12,13,45^, and thus offspring loss could influence maternal social behavior. In other words, yearly or lifetime measures of maternal sociality could themselves be influenced by offspring survival, introducing circularity in predictor and outcome variables^13^. Second, measures of sociality over a female’s lifetime may not be appropriate because sociality may change considerably as a female ages^33,46,47^; see below).

We used model comparisons to test whether female social integration was associated with offspring survival to age 1, the period of highest mortality among chimpanzees before late adulthood^33,48–50^, and when infants depend almost entirely on their mothers’ milk for their nutritional needs^51^. Using a dataset of 110 offspring born to 37 unique mothers, we constructed a null model with a binary outcome variable indicating offspring survival to age 1 and terms for offspring sex and the count of sexually mature females in the community as predictor terms (see STAR methods for list of full model terms that were removed during model selection, including maternal rank score). We then used information theoretic model comparisons of models including terms from the null model plus measures of female social integration to determine if they improved model fit. Two models had lower AICc scores than the Null model (Table 1). The best model (ΔAICc = -4.033 relative to the null model) accounted for 64% of total model weight and included the Composite Sociality Index with Females (CSI-F, a composite measure of association and grooming rates with other females). The next best fitting model (ΔAICc = -2.031 relative to the null model) accounted for 23% of total model weight, and included both CSI-F and the CSI-M (a composite measure of association and grooming rates with males). The null model was within 5 AICc points of the best-fitting model, so also received some support (Akaike weight = 0.085).

**Table 1:**
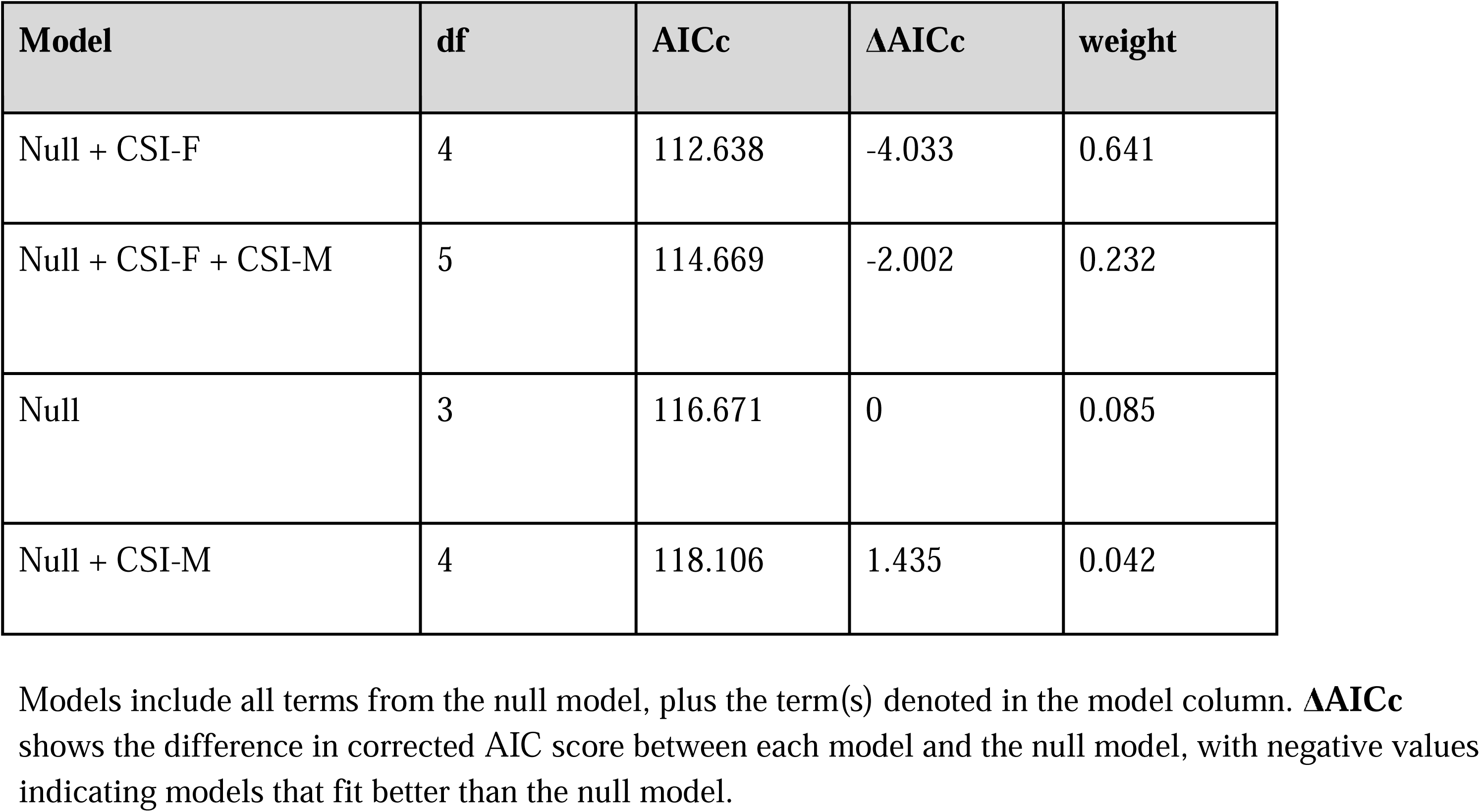
List of models and model fit parameters using measures of social integration to predict offspring survival age 1.

In the best model, social integration with females was positively associated with offspring survival to age 1, such that the odds of offspring survival to one year increased by a factor of 3.41 (95% CI 1.27 – 11.38) with a one-unit increase in maternal CSI-F in the year preceding birth (Figure 1A). In other words, holding other predictors at their mean values, a female with a CSI-F value of 0.5 (half the mean value in the year preceding the birth of her offspring) had a 74.9% probability of her offspring surviving its first year, whereas a female with a CSI-F value of 2 (twice the mean value in the year preceding the birth of her offspring) had a 95.0% probability of offspring survival to age 1 (Figure 1A; full model results: Supplementary Material).

**Figure 1:**
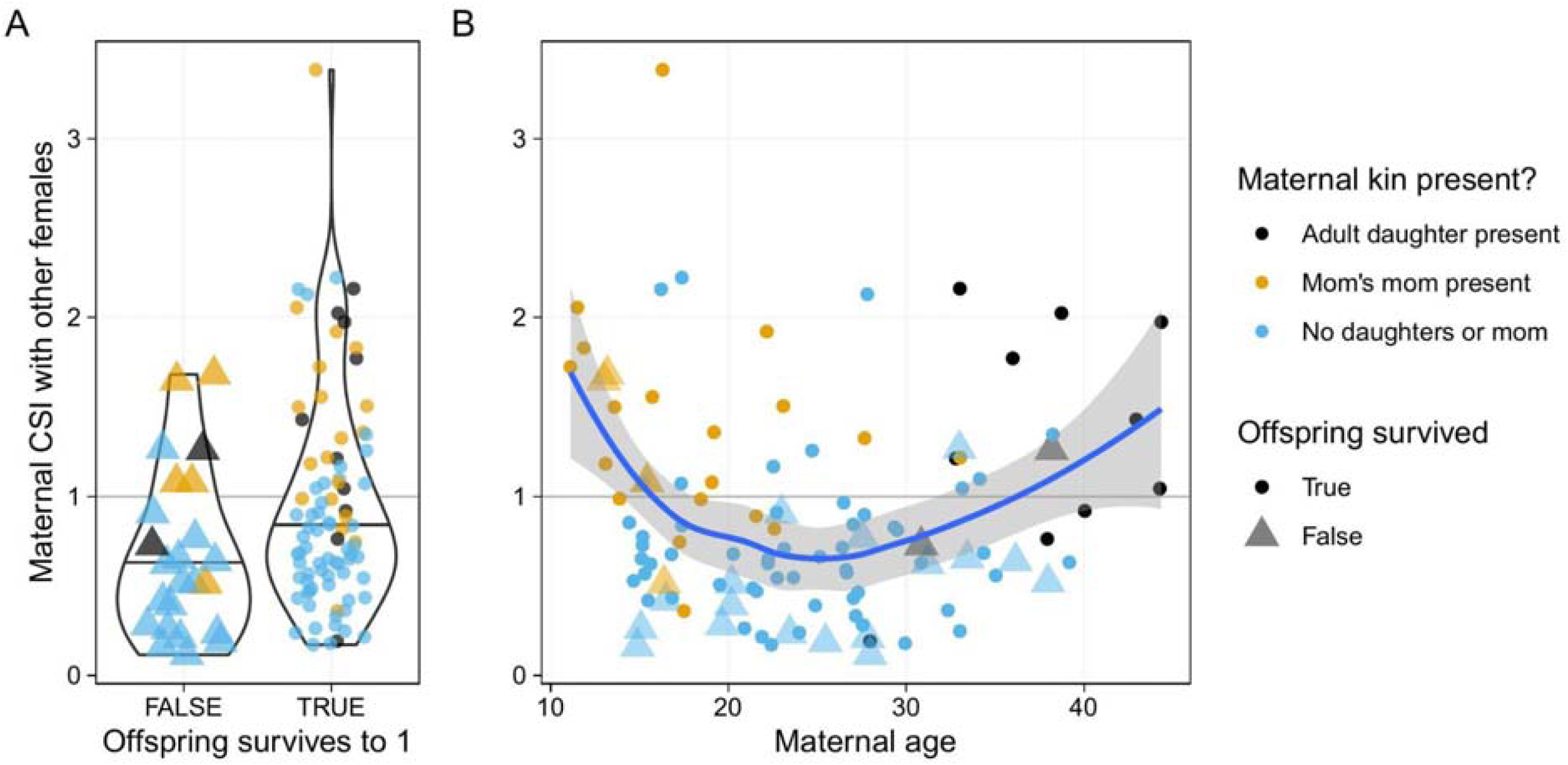
Observed values of CSI-F vs kin presence and A) offspring survival and B) maternal age. Triangles indicate offspring that died before age 1, and the blue line in B) is a loess curve drawn through the data. Points in A) are shown with jitter (in the x direction only) to better show their distributions.

Social integration with males (CSI-M) did not appear in the best-fit model, and although it did so in the second-best model, CSI-M was not associated with offspring survival in this model (OR = 1.18, CI = [0.53 – 2.79]). Furthermore, the model including only CSI-M (and terms from the Null model) had a higher AICc score than the Null model (Table 1). Additional controls indicated that results could not be explained by differences in individual observation time among mothers in the sample, and that the CSI-F measure better captured social dynamics associated with offspring mortality than did its component parts (see Supplementary Material). Together, these results suggest that a female’s social integration with males was not an important predictor of offspring survival to age 1.

We then repeated our modeling procedure for offspring survival to age 5 (the approximate age of weaning in Gombe^52–55^), instead of age 1 (see Supplementary Material). The data set for these models differed slightly, and the null model included only infant sex, but overall results remained consistent. The best model included the term from the null model plus CSI-F (ΔAICc relative to the Null model = -5.56, model weight = 0.530), and the next best model included both CSI-F and CSI-M (ΔAICc = -4.82, model weight = 0.366). CSI-F was positively associated with offspring survival in the best model, (OR = 2.42, CI = [1.01 – 6.64]) and the second-best model (OR = 2.45, CI = [1.02 – 6.75]), while in the second-best fitting model CSI-M was not associated with offspring survival (OR = 1.56, CI = [0.70 – 3.64]). Thus female social integration with other females was positively associated with offspring survival through the period of highest infant mortality and to weaning age, even when excluding mothers that died in those periods.

This analysis followed studies of sociality and offspring survival in other species^9^ in investigating the consequences of social integration (i.e. females’ overall affiliative tendency^5^, here measured with the CSI). However, different measures of sociality have been found to predict fitness-related outcomes across different studies, with no obvious way to predict a priori which measure is best^5,56^. We therefore conducted a supplementary analysis to compare the explanatory power of social integration with social bond strength (or “bondedness”^6^; i.e. the strength of an individual’s top affiliative social bonds, another common approach to measuring sociality^5^). We found that social integration (CSI-F) was the best predictor of offspring survival, although a measure of the strength of each females’ strongest social bond with other females had nearly as strong support as a predictor of offspring survival (Table S4).

### Does kinship explain the link between sociality and offspring survival?

Given the association between maternal social integration with females and offspring survival, we explored the predictors of inter-individual heterogeneity in female social integration. We found that target females’ bond strength to their top bond partner was considerably stronger if the top bond partner was close maternal kin (Supplementary Material).

We next fit a path model using our survival to age 1 dataset to investigate whether kinship explained the link between sociality and offspring survival. Path analysis revealed that females had higher CSI-F values if they had close female maternal kin in the community, and that young and old females had stronger CSI-F values; however, only CSI-F in the year preceding birth was directly associated with offspring survival to age 1 (Table 2, Figure S3). Thus kin presence was *indirectly*, but not directly, associated with offspring survival via its association with female social integration. We repeated this analysis with the survival to age 5 dataset, and found similar results (Supplementary Material).

**Table 2:** Results from path analysis of sociality and offspring survival to age 1, representing a hypothesized set of causal relationships among the maternal social environment, maternal social integration, maternal characteristics, and offspring survival (visualized in Fig. 2). Estimates represent unstandardized path coefficients, and effects whose 95% confidence intervals exclude 0 are bolded.

| Term | Estimate (CI) | p-value |
| --- | --- | --- |
| CSI-F ~ |  |  |
| **Maternal female kin** | **0.47 (0.28 – 0.66)** | **< 0.001** |
| **Maternal age** | **-0.20 (-0.36 – -0.04)** | **0.01** |
| **Maternal age<sup>2</sup>** | **0.10 (0.01 – 0.19)** | **0.02** |
| Maternal Elo | 0.14 (-0.01 – 0.29) | 0.07 |
| Maternal Elo ~ |  |  |
| Maternal female kin | 0.24 (-0.03 – 0.51) | 0.08 |
| **Maternal age** | **0.80 (0.66 – 0.94)** | **< 0.001** |
| Survival to 1 ~ |  |  |
| **CSI-F** | **0.83 (0.23 – 1.44)** | **0.01** |
| Maternal female kin | -0.32 (-0.98 – 0.35) | 0.35 |
| Maternal age | 0.05 (-0.40 – 0.50) | 0.84 |
| Maternal age<sup>2</sup> | -0.11 (-0.4 – 0.18) | 0.46 |
| Maternal Elo | 0.09 (-0.31 – 0.5) | 0.66 |

Finally, to more directly test whether kinship was responsible for the link between sociality and offspring survival, we excluded females that had female maternal kin in the community when they gave birth and re-ran our best-fitting model. Of the original 110 offspring in the sample, 40 (born to 16 females) were excluded, leaving a dataset of 70 offspring born to 27 females. As in the full data set, social integration with females was positively associated with offspring survival to age 1, such that the odds of offspring survival to one year increased by a factor of 17.27 (95% CI 1.80 – 288.08) with a one-unit increase in maternal CSI-F in the year preceding birth (Figure 2). More socially-integrated females had higher offspring survival even among the substantial majority of females in the community that lacked maternal female kin.

**Figure 2:**
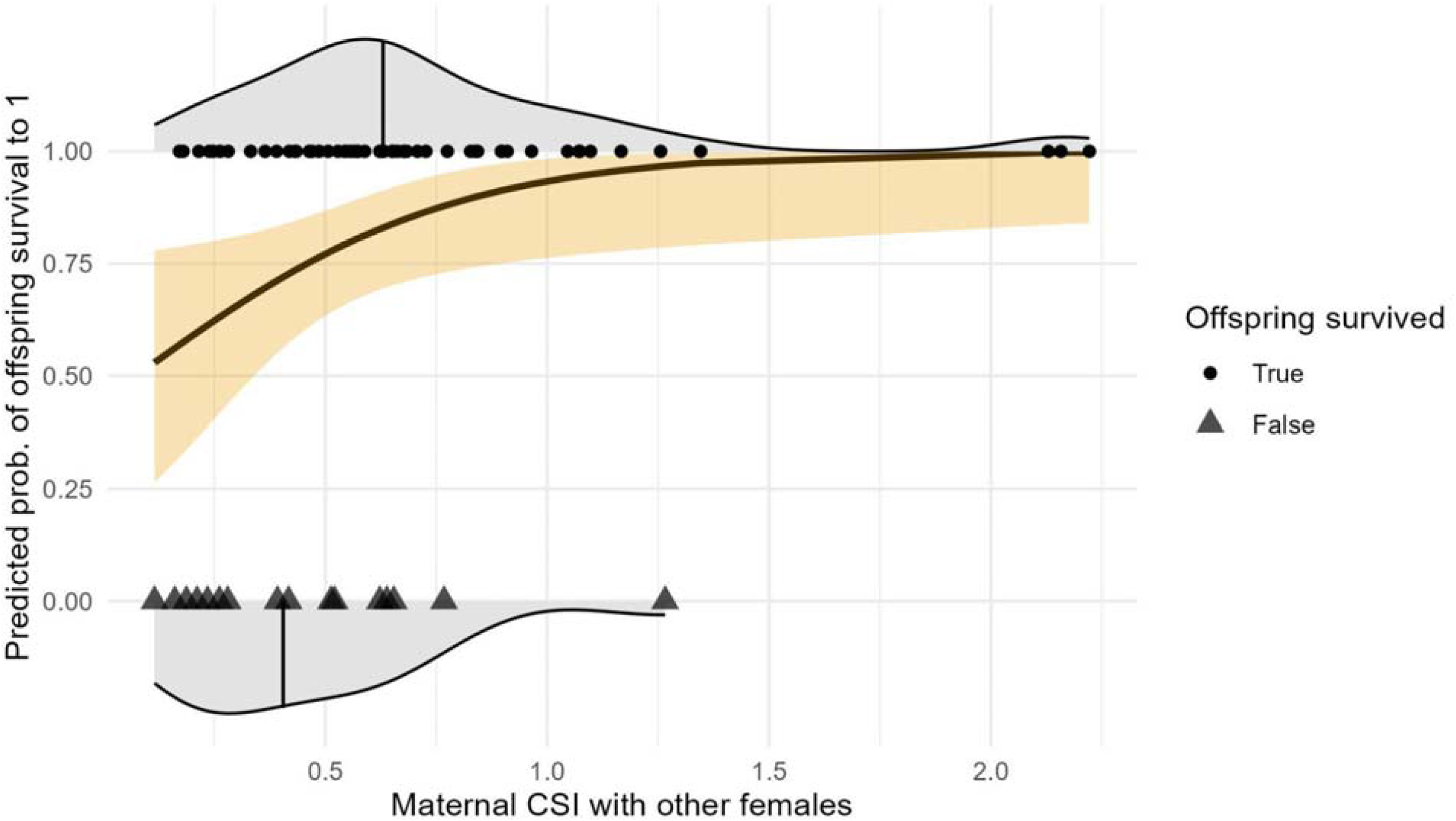
Predicted probability of offspring survival vs. maternal CSI-F for the 70 offspring born to females without female kin in the community at birth. Points and top density plot show the empirical distribution of maternal CSI-F for those offspring that survived to age 1, while triangles and bottom density plot indicate the empirical distribution of maternal CSI-F for those offspring that died before age 1. The vertical line in each distribution shows the median maternal CSI-F value for offspring that survived and those that died, respectively.

### Maternal sociality before and after birth

Prenatal social and ecological conditions can influence offspring health and survival directly^57–59^, and/or be relevant to offspring survival if they are predictive of conditions after birth. To determine whether maternal sociality in the year before birth was predictive of maternal sociality after birth, we restricted our dataset to offspring that survived their first year, and whose first year of life did not extend beyond the end of the study period (81 offspring born to 36 females). This is because infant death may have unpredictable effects on maternal social behavior. Among females whose offspring survived the first year, CSI-F values in the year preceding birth were strong predictors of CSI-F values in the year after birth (Pearson correlation = 0.580), and in the regression model CSI-F (before birth) was significantly positively associated with CSI-F (after birth) (□ = 0.45, 95% CI = [0.19 – 0.71], *p* < 0.001). The presence of female kin in the year after birth was also associated with higher CSI-F values in the same period (□ = 0.60, 95% CI = [0.28 – 0.92], *p* < 0.001).

In contrast, CSI-M in the year preceding birth was a weaker predictor of CSI-M in the year after birth among females whose offspring survived the first year (Pearson correlation = 0.303); in the regression model CSI-M (before birth) was again not as strongly associated with CSI-M (after birth) (□ = 0.24, 95% CI = [-0.02 – 0.50], *p* = 0.072) as were CSI-F pre- and post-birth. Similarly, the presence of male kin in the year after birth was not associated with higher CSI-M values in the same period (□ = 0.21, 95% CI = [-0.17 – 0.59], *p* = 0.267).

Thus, maternal social integration with females was consistent across the two-year period surrounding birth, whereas maternal social integration with males was not. The former result may hold for even longer periods, as evidence from western chimpanzees (*P. t. verus*) suggests that interindividual differences in female grooming and affiliative tendencies are repeatable over periods of three to 11 years ^60^.

## Discussion

Female chimpanzees that were more socially integrated with other females in the year preceding the birth of an offspring had higher offspring survival to age 1 and to age 5. Furthermore, among females whose offspring survived to age 1, social integration with other females in the year preceding birth was a strong predictor of social integration with other females in the year after birth, suggesting a consistency of social environment across these periods. Although females formed strong bonds with adult female kin, and those with female kin in the community were more strongly socially integrated with females overall, kin presence did not directly predict offspring survival. Indeed, most females in the study did not have access to close maternal female kin, and among this subset social integration remained strongly predictive of offspring survival through the critical life stages assessed here. Furthermore, we have shown that female social integration is an important predictor of offspring survival in a species with 1) high fission-fusion dynamics and 2) a non-matrilineal dominance hierarchy, underscoring the importance of studies of female sociality in a wide array of social organizations.

In contrast, female social integration with males was not associated with offspring survival. Furthermore, the male social environment was less consistent across pre- and post-birth periods. In certain baboon species (e.g. olive, yellow, chacma), male friendships with females are associated with paternity, and with male protection from harassment and infanticide after birth^24,61,62^. In chimpanzees, however, despite evidence that males bias some affiliative behaviors towards their offspring^63,64^, it is not known whether males selectively protect or intervene on behalf of their offspring. And although females in Gombe associate disproportionately with adult brothers and sons after giving birth^65^, whether these associations provide protection against infanticide remains unexplored.

Although an important contributor to infant mortality^66^, maternal mortality cannot explain current results because we excluded offspring of females that died shortly after giving birth. In addition, in chimpanzees and several other primate species, offspring of mothers that die within four years of giving birth have higher mortality even before their mothers’ death^67^. Yet our analysis of offspring survival to age five excluded offspring whose mothers died within five years of giving birth, and the association between maternal sociality and offspring survival remained. Impending maternal death cannot then explain the results presented here.

As in studies of female cercopithecine monkeys^3,12^, the association between sociality and adaptive outcomes described here was not mediated by dominance rank. Maternal Elo score was not associated with offspring survival to age 1, either directly or indirectly via an association with female social integration. Using a restricted dataset to model offspring survival to age 5, maternal Elo score was *indirectly* (but not directly) associated with offspring survival via its association with female social integration. Previous analyses in Gombe found that rank predicted offspring survival to age 7^31^ and to age 8^68^. Why this discrepancy? Supplementary analysis reveals that it likely results from a changing association between maternal rank and offspring survival over decades; high-ranking females had considerably higher offspring survival to age 7 than did low-ranking mothers in the 1970’s, whereas offspring survival did not differ by maternal rank score in the current study period (Supplementary Material, Figure S5). Follow-up work is required to determine the cause of this changing association between maternal rank score and offspring survival.

Unlike in capuchins and cercopithecines, in which previous evidence of a positive association between sociality and offspring survival is reported^9,10,12,20^, kinship among the present population of female chimpanzees is not the primary determinant of social structure. The majority of offspring in our sample (70 of 110) were born to mothers that did not have adult female maternal kin in the community, and social integration with females remained an important predictor of offspring survival among the cohort of females without access to close female kin.

What mechanisms might link female social integration in the year preceding birth to offspring survival? Given the consistency of maternal social integration with other females pre- and post-birth, one possibility is that social integration could facilitate tolerance from other females during the prenatal period and/or first year of life. This could promote maternal feeding efficiency or reduce received aggression, including infanticide in the postpartum period^5,6^. Previous studies have estimated a minimum of 3.5% and maximum of 19% mortality in the first year due to female-led infanticide in our study population^69^, although we currently lack data to resolve the relationship between social integration and infanticide in the current analysis.

Additionally, while competitive interactions among females do not tend to influence relative rank in the Gombe population^43^, females still compete aggressively for access to food resources^70^. Female chimpanzees in Kanyawara, Kibale National Park, Uganda, formed more aggressive coalitions with other females with whom they had a social bond, but coalition formation was no more likely for kin or strongly-bonded dyads than for weakly-bonded dyads^71^. Thus, more socially integrated females could receive more agonistic support if they form more (but not necessarily stronger) social bonds, which could provide protection from harassment or facilitate joint defense of food patches^5,6^. Follow-up research should therefore explore the behavioral correlates of strong social integration with females. Additionally, work to study the influence of social integration on other components of female fitness could help to resolve the mechanisms responsible for the result reported here.

Alternatively, both female social integration and offspring survival could depend on a third factor like individual quality (e.g. refs. ^72–74^). In that case, females in good condition, or those with advantageous genotypes, may invest more both in forming social bonds and in offspring care. In Gombe, higher body mass is associated with shorter inter-birth intervals, suggesting that body size could be a proxy for individual quality^73^, although it is not known whether these measures are related to female social behavior. However, if interindividual variation in female quality was responsible for the results presented here, females should differ in their abilities to raise offspring to age 1 or age 5 over repeated reproductive events; instead, in the current analysis, maternal identity did not account for any variance in offspring survival. Furthermore, studies in humans find that the positive association between social ties and fitness-related outcomes, like health and survival, remain even after controlling for initial health, socioeconomic status, and other factors that might jointly influence sociality and fitness outcomes^2,75^, suggesting an independent effect of sociality on adaptive outcomes.

If female chimpanzees form stronger bonds with female kin, and appear to benefit from being socially integrated, why do chimpanzees show a derived pattern of female dispersal^76^? Models of primate socioecology have tended to emphasize the mode of female competition over resources in explaining dispersal patterns, with female dispersal arising when females do not benefit from cooperative resource defense^77–80^. An alternative proposal involves local resource enhancement^76^; if males gain sufficient benefits from remaining in their natal group and cooperating with kin (or developing cooperative relationships with non-kin over time^81^), females could be forced to emigrate to avoid inbreeding^76,82^. Although the mechanism requires further investigation, the current results, along with recent evidence that males gain similar reproductive benefits from strong social bonds^83^, are more consistent with the latter scenario; in chimpanzees, both males and females that are more socially integrated have higher reproductive success. But in chimpanzees, overlapping female ranges may provide males with unusually high benefits from remaining with kin and cohort mates to cooperatively defend a group territory^84^, forcing females to emigrate to avoid inbreeding^76^.

But like humans in patrilocal societies, where women nevertheless form strong social networks with non-kin^85–87^, female chimpanzees manage to form strong social bonds even with non-kin^34,38^. In the current analysis offspring survival did not differ as a function of dispersal status. Instead, in our closest relatives, offspring survival is associated with female social integration, in the absence of female philopatry and the strict matrilineal kinship structure seen in many other haplorhine primates.

### Limitations of the study

As with any observational study, effects reported here are correlational, and further work is required to establish the causal relationship between female social integration and offspring survival (see above discussion of potential mechanisms). Furthermore, correlations between a behavior and adaptive outcomes can change over time, as demonstrated in the Supplementary analysis of the association between female dominance rank and offspring survival before and during the current study period. Such changes could be due to changing environmental conditions (e.g. ref. ^88^), a changing social environment (e.g. ref. ^89^), or for other reasons. In a long-lived and socially variable species like chimpanzees, data from other populations, and additional decades of research, are required to establish the universality of the results reported here.

## Acknowledgements

We thank Susan Alberts, Beniamino Tuliozi, Brian Lerch, Maria Creighton, and three anonymous reviewers, for helpful comments on the analysis and manuscript, and the wonderful data collection team at Gombe for their tireless and careful work. Data collection was supported by the Jane Goodall Institute, construction of the long-term database was supported by grants from the NSF (DBS-9021946, SBR-9319909, BCS-0452315, IOS-1052693, IOS-1457260), the Harris Steel Group, the Windibrow Foundation, the University of Minnesota, and Duke University.

## Author contributions

Conceptualization, JTF, KKW, AEP; Formal Analysis, JTF; Data Curation, AEP, EVL, DM, JTF; Writing – Original Draft, JTF, KKW, MAS, EVL, CMM, DM & AEP; Writing – Review & Editing, KKW, MAS, EVL, CMM, DM, & AEP; Visualization, JTF; Funding Acquisition, AEP.

## STAR Methods

### Key Resources Table

STAR Methods Key Resources Table
| REAGENT or RESOURCE | SOURCE | IDENTIFIER |
| --- | --- | --- |
| Deposited Data |  |  |
| Cleaned data and code | https://github.com/jtfeld/fem_soc_offspring_surv | Zenodo: DOI [10.5281/zenodo.10070424](https://doi.org/10.5281/zenodo.10070424) |
| Experimental Models: Organisms/Strains |  |  |
| 110 offspring and 37 adult female chimpanzees | Not applicable | Not applicable |
| Software and Algorithms |  |  |
| R 4.3.1 | R Core Team, 2023 | https://cran.r-project.org/ |
| R package EloOptimized 0.3.1 | Feldblum et al., 2023 | https://cran.r-project.org/web/packages/EloOptimized/index.html |
| R package lme4 1.1-35 | Bates et al., 2015 | https://cran.r-project.org/web/packages/lme4/index.html |
| R package lavaan 0.6-16 | Rosseel 2012 | https://cran.r-project.org/web/packages/lavaan/index.html |
| R package lavaanPlot 0.8.1 | Lishinski 2023 | https://cran.r-project.org/web/packages/lavaanPlot/index.html |
| R package MuMIn 1.47.5 | Bartoń 2023 | https://cran.r-project.org/web/packages/MuMIn/index.html |

### Resource Availability

#### Lead contact

Further information and requests for code and data should be directed to and will be fulfilled by the lead contact, Joseph T. Feldblum.

#### Materials availability

This study did not generate new unique reagents.

#### Data and code availability

- Data are available at https://github.com/jtfeld/fem_soc_offspring_surv
- All original code is available at https://github.com/jtfeld/fem_soc_offspring_surv, and permanently archived at https://zenodo.org/records/10070425 (DOI: 10.5281/zenodo.10070424) and is publicly available as of the date of submission.
- Any additional information required to reanalyze the data reported in this paper is available from the lead contact upon request.

### Experimental model and subject details

#### Subjects

Subjects were 110 offspring and 37 adult female free-ranging chimpanzees (*Pan troglodytes schweinfurthii*) in the Kasekela community in Gombe National Park, Tanzania. Females were between 11.1 to 44.4 years old, with a median age of 23.0 years (see below for inclusion criteria).

#### Ethical guidelines

Data collection was approved by Tanzania National Parks, Tanzania Wildlife Research Institute, and Tanzania Commission for Science and Technology, as well as the Duke University Institutional Animal Care and Use Committee.

### Method details

Data come from long-term observations of free-ranging eastern chimpanzees in Gombe National Park, Tanzania. Researchers have conducted daily focal follows of individually-recognized chimpanzees in the Kasekela community since 1968, one of three communities in the park^33,90^. Chimpanzees have high fission-fusion dynamics^30^, in which the full community rarely forms into a single party and party composition changes frequently^33^. Thus chimpanzees have some measure of control over their association partners, and patterns of association should reflect partner preference^35,83,91^. Observers record party composition at 15-minute intervals, as well as the time of arrival and departure for every individual joining and leaving the focal individual’s party. Observers also record all occurrences of social behaviors, including focal grooming, aggressive interactions, and submissive vocalizations.

Researchers in Gombe have been collecting fecal and urine samples to identify SIVcpz infection using noninvasive measures since 2000. Briefly, samples are tested for virus-specific antibodies by western blot analysis, and infection is confirmed by RT-PCR analysis^92–94^.

The current sample includes offspring born to mothers of known dominance rank in the Kasekela community, during the period for which we have data on grooming and dominance rank. We began with a dataset of 113 offspring born to 38 mothers between 1980 and 2017.

We first analyzed factors influencing offspring survival to age 1, as this is the period of highest mortality among chimpanzees before late adulthood^33,48–50^. Because chimpanzees younger than age 1 invariably die if their mother dies^66,95^, we excluded two offspring whose mother died before they would have turned 1. We also excluded one offspring born to a peripheral female (KP) who was never observed associating with males or females in the year prior to birth. Because of this, we were unable to generate an index of her sociality, as her grooming rate was undefined (see below). However, this offspring died before age 1, so this data point is consistent with the main result reported here. These exclusions resulted in a data set of 110 offspring born to 37 mothers. Mothers in the sample ranged from 11.1 to 44.4 years old, with a median age of 23.0 years. 24 of 110 offspring in the sample died before age 1 which is consistent with early life mortality patterns reported previously^50^.

Of the 110 offspring in the sample, 51 were born to 23 immigrant mothers, 53 were born to 12 natal mothers, and 6 were born to 2 females of unknown residence status. Natal and immigrant mothers differed considerably in their presence of close maternal female kin (hereafter “female kin”) in the community. Among offspring born to immigrant mothers, 8 of 51 were born to mothers with female kin in the community in the year preceding birth. This figure was 31 of 53 for offspring born to natal mothers, and 1 of 6 offspring born to mothers of unknown residence status (See Table S1 for these data broken down by kin class.). This variation allowed us to investigate the importance of kinship in facilitating the fitness effects of maternal sociality.

We then repeated this analysis for offspring survival until age 5, instead of age 1, for two reasons. First, we conducted this second analysis to facilitate comparison with earlier studies of maternal sociality and reproductive success that modeled the production of weaned offspring (e.g. refs. ^96,97^); chimpanzee offspring are typically weaned around age 5^52–55^. Second, we wanted to reduce the potential influence of impending maternal death on offspring mortality in our analysis. A recent study reported that, in chimpanzees and several other primate species, impending maternal death, or maternal death within four years after giving birth, was associated with higher offspring mortality even if mothers outlived their offspring^67^. In the analysis of survival to age 5, we excluded offspring whose mothers died within 5 years of giving birth, thus excluding offspring born to mothers who died during the critical window described in ref. ^67^. Despite these changes, results were nearly identical to those reported in the main text (see Supplementary Material for full description of the analysis and results).

### Dominance rank data

Following previous work in Gombe^43^ we generated dominance rank scores for females using submissive pant-grunt vocalizations between females with clearly identified actor and recipient, as well as aggressive interactions between females with clearly determined winner and loser. Because the youngest mother in the sample was 11.1 years of age at birth, we included pant grunts and decided aggression events where both actor and recipient were at least 11 years old. This resulted in a sample of 1384 agonistic interactions among 50 females, recorded between 1969 and 2018. Females in the current analysis had a median of 36 agonistic interactions (range 4 to 308).

To generate rank scores, we used a modified version of the Elo rating method^98^ that uses maximum likelihood estimation to fit the scaling parameter and initial Elo scores^43^, implemented in the EloOptimized package in R (version 0.3.1; ref. ^99^).

### Other covariates

We included data on several other predictor terms that 1) have been shown in previous analysis to be associated with mortality in chimpanzees, or 2) may be confounds for a possible relationship between sociality and infant mortality. First, we included data on maternal SIV status at birth. SIV infection was associated with increased infant mortality in an earlier analysis in Gombe^92^. Only 7 offspring in the sample were born to SIV positive mothers, 92 to SIV negative mothers, and 11 to mothers with unknown SIV status. Following earlier analyses in Gombe^73^, we coded SIV positive females as 1, SIV negative females as -1, and females of unknown SIV status as 0. This approach maximizes sample size while avoiding introducing bias in parameter estimates^73^.

We additionally included data on maternal immigrant status, as immigrant females rarely co-reside with kin, and may thus be expected to be less social. If immigrants differ from natal females in their sociality, and also differ from natal females in their rate of offspring survival due to resource availability or other factors, immigrant status could be responsible for an association between sociality and offspring survival. As above, we coded immigrant females as 1, natal females as -1, and females of unknown immigrant status as 0 to maximize sample size.

In Gombe, maternal age was associated with offspring survival in one study ^68^ but not others^100,101^. Since associations between offspring survival and maternal parity, age and age^2^ have also been documented in other primates^101^ we included these terms in our full models. To ensure comparable effect sizes, we standardized maternal age by Z transformation across the entire dataset, and included a squared standardized age term to account for potential differences in offspring survival among young or old mothers. 24 offspring in the data set were primiparous births, 84 were born to parous mothers, and two were of unknown firstborn status. As above, we coded first-born offspring as 1, non-first-born offspring as -1, and offspring of unknown first-born status as 0.

To account for the possibility of increased risks of infanticide or feeding competition in bigger groups, we included measures of the count of males over age 12 and females over age 11 in the community on each offspring’s date of birth.

Finally, we included a measure of offspring sex among the candidate predictor terms in our full model; an analysis of mortality data from five study populations found that mortality rates differed between males and females throughout the lifespan^48^, although this is not always the case among young chimpanzees^100,102–104^. The data set included 60 male offspring, 42 female offspring, and 8 offspring of unknown sex, which we coded as 1, -1, and 0, respectively.

### Sociality measures

We created individual-level composite sociality indices (hereafter “CSIs”) to measure female social integration in the year preceding her offspring’s birth date^9^. We measured sociality over a year to facilitate comparison with previous analyses of sociality and adaptive outcomes in chimpanzees and other non-human primates^10,14,15,56,83,91^.

As noted above, we used data from the year before birth to predict offspring survival to avoid introducing circularity in predictor and outcome variables (see Results for full discussion). Singleton pregnancies at Gombe last a mean of 228 days^102^, so the year before birth includes periods of both reproductive cycling and pregnancy.

We included females as potential social partners if they were at least 11 years of age and still alive on the date of the target female giving birth; the youngest female in our dataset was 11.1 years of age when she gave birth, and 11.5 is the average age of sexual maturity for females in Gombe^44^. We included males as potential social partners if they were at least 13 years of age and still alive on the date of the target female giving birth, to ensure that they were at least 12 years of age by the start of the window. Around age 12, male chimpanzees begin to travel independently from their mothers, and begin to compete aggressively for access to females^105,106^.

To measure female social integration, we calculated CSIs based on female association time and grooming rate with community members. We included two behavioral measures in CSI scores: 1) summed time females were observed in the same party with community members, and 2) the rate at which females groomed with other community members.

We first summed the time each target female was observed with other community members in the year before giving birth. We then divided each female’s summed time with community members by the mean value of summed time with community members among all females in the community during the 1-year window.

We repeated this process to calculate each female’s scaled grooming rate. We summed the minutes each female groomed with others (any direction), and divided that by the summed focal time each female spent with others. We then divided each female’s overall grooming rate by the mean overall grooming rate among all females in the community during the 1-year window.

These measures could thus take values in [0, ∞), with values above 1 indicating that a female associated or groomed with others above the mean rate for females in her community during the previous year, and those below 1 indicating that a female associated or groomed with others below the mean rate during the previous year. Results based on this method of standardizing sociality measures by dividing by the mean value in each window were substantively similar to those based on standardizing by Z transformation, and models based on the method presented here fit slightly better.

We calculated each female’s CSI separately with males and females. The CSI-F (CSI with females) thus represented female social integration with other females, relative to that of other females in the same period. Similarly, the CSI-M (CSI with males) represented female social integration with males, relative to that of other females in the same period. Our analysis thus asked whether females that associated with and groomed others at high rates *relative to other females in her community* in the year preceding the birth of a given offspring were more or less likely to have their offspring survive their first year of life.

We also explored standardizing association time and grooming rates by Z transformation, instead of by dividing by the mean value in each window. Model results and model comparisons were substantively similar, but models fit slightly better using the ratio method described above, so we report models here using the ratio method.

### Statistical analysis of offspring survival

We used logistic regressions to model the odds ratios of offspring survival to age 1^107^, using the lme4 and stats packages in R^108,109^. Following recent work with male chimpanzees^83^, we first constructed a global model containing all non-sociality predictor terms (offspring sex, firstborn status, maternal rank score, maternal age and squared maternal age, maternal immigration status, maternal SIV status, and counts of males and females in the community). We also explored including a random intercept term for maternal identity, but this term did not account for any variance so we excluded it from future models.

We then used a model comparison procedure, with AIC corrected for small sample sizes (AICc) as the selection criterion, to determine the best set of non-sociality terms for predicting offspring survival to age 1. We used the dredge function in the R package MuMIn^110^ to compare all subsets of the global model, excluding only models from the candidate set that included the squared maternal age term but *not* the corresponding first order term.

Because adding a completely random “nuisance” parameter to a model tends to increase AIC scores by ∼ 2 AIC points, potentially leading to inclusion in the top model set^111^, we used the nesting rule to exclude models from the top model set that differed from a better-supported model by the addition of a single term^112,113^. We considered models in the candidate set of best models if they differed from the best-supported model by less than 6 AICc points^112^. The best fitting non-sociality model included two terms: offspring sex (with male offspring less likely to survive than females; Odds Ratio: 0.66, 95% Confidence Interval: 0.38 – 1.08) and the count of adult females in the community on an offspring’s date of birth (with lower likelihood of survival when the community was larger; OR: 0.91, 95% CI: 0.81 – 1.02). We considered this model our null model against which to compare models that included measures of sociality.

We then created three test models by adding CSI-F, CSI-M, and both CSI-F and CSI-M to the null model in turn, and compared model fit to that of the null model using AICc values. Thus, the model comparison procedure sought to determine whether including a given measure of female sociality increased the likelihood of minimizing information loss when predicting offspring survival relative to the model that did not include a sociality predictor.

We ran two additional analyses to rule out an alternative explanation for the relationship between CSI-F and offspring survival reported in the main text and the appropriateness of our measure of female social integration. We first showed that the main result could not be explained by differences in female observation time, and we second showed that that composite sociality index appears to better capture social dynamics associated with offspring mortality than do its component parts (see Supplementary Material).

In addition, we repeated our modeling procedure for offspring survival to age 5, instead of age 1 (see Supplementary Material). The data set for these models differed slightly, as the null model included only offspring sex. Nevertheless, overall results were consistent, with CSI-F again present in the best model for predicting offspring survival to age 5.

### Does kinship explain the link between sociality and offspring survival?

Given the association between maternal social integration with females and offspring survival (see Results), we explored the predictors of inter-individual heterogeneity in female social integration.

Inspection of the raw data suggested that females with close adult female maternal kin in the community also had stronger CSI-F values (Fig. 1). Additionally, the youngest and oldest females tended to have particularly strong CSI-F values (Fig. 1). Therefore, to further explore the predictors of female sociality, we modeled the predictors of each female’s top dyadic bond strength and found that females’ top bond strength was higher if that bond was with their mothers, their adult daughters, or their adult maternal sisters (Supplementary Material, Figure S2). On the other hand, although CSI-F values varied predictably with age, top bond strength was not associated with maternal age or Elo score (Supplementary Material).

Finally, to explore the complex relationships between our predictor terms, CSI-F, and offspring survival, we fit a path model using our survival to age 1 dataset. As before, this data set included 110 offspring born to 37 mothers between 1980 and 2017. We conducted the analysis using the lavaan package in R^114^, using the default diagonally-weighted least squares estimator.

We specified kin presence (i.e. the presence of a female’s mother, daughter, or maternal sister over age 11), female age, and squared female age as predictor (i.e. exogenous) variables, with female Elo score as an intermediate outcome (i.e. endogenous) variable predicted by kin presence and female age, CSI-F as a second intermediate (i.e. endogenous) variable predicted by kin presence, female age, squared female age, and female Elo score, and offspring survival as a binary outcome (i.e. endogenous) variable predicted by all five other terms (Table 2; Figure S3). All tests indicated satisfactory model fit (Model □^2^ = 0.228, Df = 1, *p* = 0.633; RMSEA = 0.000, *p* = 0.677; CFI = 1.00; SRMR = 0.000). We then used the lavaanPlot package in R^115^ to visualize the associated path diagram (Figure S3).

As a final test of the relative importance of kin presence and social integration, we ran a model comparison procedure comparing our best-fit model for predicting offspring survival to age 1 to two models in which we replaced the CSI-F term with a term indicating the presence of female kin. We reasoned that, if the relationship between female social integration and offspring survival was attributable to the presence of kin, the latter model would fit better than the former. In one model we included a term indicating the presence of at least 1 female kin in the year before birth, and in the other we included a term indicating the presence of at least 1 female kin in the year after birth.

### Maternal sociality before and after birth

To determine whether maternal sociality in the year before birth was predictive of maternal sociality after birth, we restricted our dataset to offspring that survived their first year, and whose first year of life did not extend beyond the end of the study period (81 offspring born to 36 females). This is because, as noted above, infant death may have unpredictable effects on maternal social behavior. We first calculated a simple Pearson correlation between pre- and post-birth CSI-F measures. We then ran a linear mixed model using the lme4 package in R^108^ with CSI-F in the year after birth as the outcome variable, CSI-F in the year before birth and maternal kin presence in the year after birth as predictor terms, and a random intercept term for female identity.

We then repeated this analysis with CSI-M values before and after birth, including a measure of male kin presence instead of female kin presence. We considered a female to have male kin in the group if she had a brother or son that was 1) older than age 13 when she gave birth, and 2) alive throughout the year after giving birth.

### Supervising bodies

We thank Jane Goodall for permission to work with the long-term data and the Tanzania National Parks, Tanzania Wildlife Research Institute, and Tanzania Commission for Science and Technology for permission to work in Gombe National Park. All data collection was approved by these Tanzanian bodies and the Duke University Institutional Animal Care and Use Committee.

