## Supplementary Material for "Socially integrated female chimpanzees have lower offspring mortality"

#### Supplementary analyses

##### Full details of best model of survival to age 1

The best-fitting model of offspring survival to age 1 included the two terms from the null model: 1) count of sexually mature females in the community at birth and 2) offspring sex, plus CSI-F (“Composite Sociality Index with Females”, measuring maternal social integration with females). In this model, CSI-F was positively associated with offspring survival to age 1, such that the odds of offspring survival to one year increased by a factor of 3.41 (95% CI 1.27 – 11.38) with a one-unit increase in maternal CSI-F in the year preceding birth (Figure 1A). In other words, holding other predictors at their mean values, a female with a CSI-F value of 0.5 (half the mean value in the year preceding the birth of her offspring) had a 74.9% probability of her offspring surviving its first year, whereas a female with a CSI-F value of 2 (twice the mean value in the year preceding the birth of her offspring) had a 95.0% probability of offspring survival to age 1 (Figure 1A in the main text). Similarly, this model predicted that offspring born into a community with the smallest observed number of sexually mature females (n = 12) had a 91.3% probability of survival to age 1, whereas offspring born into a community with the largest observed number of sexually mature females (n = 27) had a 72.9% probability of survival to age 1. Finally, model-predicted survival probability of female offspring was 89.2%, but just 73.6% for male offspring.

##### Additional analyses of survival to age 1

We ran two additional analyses to rule out alternative explanations for the relationship between CSI-F (the Composite Sociality Index with Females) and offspring survival reported in the main text. First, to assess the appropriateness of our composite index of sociality, we ran an additional model comparison procedure using the survival to age 1 dataset (see Methods for details). We compared the best-fitting model for predicting offspring survival to age 1 (which included the CSI-F measure) with two models in which we replaced the CSI-F measure with one of its component affiliative measures, along with our null model. The models that included scaled association with females and grooming rate with females (the two components of the CSI-F measure) both fit better than the null model (ΔAICc relative to null model = -0.54 & -2.69, respectively). However, neither fit better than the CSI-F model (ΔAICc relative to null model = -4.03). In the association time model, scaled time with other females was positively associated with offspring survival (OR = 2.62, 95% CI = 0.83 – 8.68), and in the grooming rate model, scaled grooming rate with other females was also positively associated with offspring survival to age 1 (OR = 1.95, CI = 1.07 – 4.36). However, because model fit for the CSI-F model was considerably better, the composite sociality index appears to better capture social dynamics associated with offspring mortality than do its component parts.

Second, some females in Gombe are more peripheral than others, leading to differences between females in time observed. To ensure any association between social integration and offspring survival was not just driven by differences in female observation time, we ran an additional model with a term for total female observation time (relative to the female mean) in the year before birth. This female observation time model had a marginally higher AICc score than the Null model (ΔAICc = 0.85), whereas the best-fitting model had a considerably lower AICc score than the Null model (ΔAICc = -4.03), indicating that the CSI-F result cannot be explained by differences in female observation time.

As a final test of the relative importance of kin presence and social integration, we ran a model comparison procedure comparing our best-fit model for predicting offspring survival to age 1 to two models in which we replaced the CSI-F term with a term indicating the presence of female kin. We reasoned that, if the relationship between female social integration and offspring survival was attributable to the presence of kin, the latter models would fit better than the former. In this model comparison procedure, both kin presence models had considerably higher AICc values than the CSI-F model (kin presence in the year after birth: ΔAICc = 5.63 relative to the CSI-F model; kin presence in the year before birth: ΔAICc = 5.75 relative to the CSI-F model). Thus, consistent with the results from the path analysis above, female social integration was a better predictor of offspring survival than was kin presence.

##### Survival to age 5

We repeated our modeling procedure using logistic models of offspring survival, but for survival to age 5. To do this, we further restricted our data set to offspring born to females that survived at least 5 years after giving birth. This resulted in a data set of 97 offspring born to 29 mothers from 1980 to 2017. 38 of the 97 offspring in this sample died before age 5.

We used logistic regressions to model the odds ratios of offspring survival to age 5 ^1^, using the lme4 and stats packages in R ^2,3^. As with the primary analysis, we first constructed a global model containing all non-sociality predictor terms (offspring sex, firstborn status, maternal rank score, maternal age and squared maternal age, maternal immigration status, maternal SIV status, and counts of males and females in the community). As before, the random intercept term for maternal identity did not account for any variance, so we excluded it from future models.

We again used a model comparison procedure, with AIC corrected for small sample sizes (AICc) as the selection criterion, to determine the best set of non-sociality terms for predicting offspring survival to age 5. We used the dredge function in the R package MuMIn^4^ to compare all subsets of the global model, excluding only models from the candidate set that included the squared maternal age term but *not* the corresponding first order term.

We again used the nesting rule to exclude models from the top model set that differed from a better-supported model by the addition of a single term^5,6^, and considered models in the candidate set of best models if they differed from the best-supported model by less than 6 AICc points^5^. The best fitting non-sociality model included just one term: offspring sex. We considered this model our null model against which to compare models that included measures of sociality.

We then created three test models by adding CSI-F, CSI-M, and both CSI-F and CSI-M to the null model in turn, and compared model fit to that of the null model using AICc values. Thus the model comparison procedure sought to determine whether including a given measure of female sociality increased the likelihood of minimizing information loss when predicting offspring survival relative to models including no sociality predictor.

We again also included a model with a term for total female observation time (relative to the female mean) in the year before birth. Finally, finding evidence of an association between CSI-F and offspring survival, we ran two additional models, replacing CSI-F with its component parts (scaled time with other females, and scaled grooming rate with other females). This was to determine the relative importance of each component of the CSI-F measure. We compared model fits using AICc.

Overall, results of the survival to 5 analysis were remarkably similar to those of the survival to 1 analysis reported in the main text.

Three models had lower AICc scores than the Null model (Table S2). The best model accounted for 53% of total model weight and included the Composite Sociality Index with Females (CSI-F). The next best fitting model accounted for 37% of total model weight, and included both CSI-F and the CSI-M. Finally, the third best fitting model included the term for total female observation time, but accounted for only 4% of total model weight.

In the best model, social integration with females was positively associated with offspring survival to age 5, such that the odds of offspring survival to one year increased by a factor of 3.22 (95% CI 1.38 – 8.64) with a one-unit increase in maternal CSI-F in the year preceding birth. In other words, a female with a CSI-F value twice the mean value in the year preceding the birth of her offspring was predicted to have a 3.22-fold increase in the odds of her offspring surviving its first year relative to a female with a CSI-F value at the mean.

While the female observation time model had a marginally lower AICc score than the Null model, this model accounted for just 4% of model weight, indicating, as with the primary analysis, that the CSI-F result cannot be explained by differences in female observation time.

Social integration with males (CSI-M) did not appear in the best-fit model, and although it again did so in the second best model (ΔAICc = 0.741 relative to the best model), CSI-M was not associated with offspring survival in this model (OR = 1.58, CI = [0.75 – 3.50]). Furthermore, the model including only CSI-M again had a higher AICc score than the Null model (Table S2). Together, these results suggest that female social integration with males was not an important predictor of offspring survival to age 5.

In a subsequent model comparison, models including scaled association with females and grooming rate with females (the two components of the CSI-F measure) both fit better than the null model (ΔAICc relative to null model = -0.40 & -3.85, respectively). However, as before, neither fit better than the CSI-F model (ΔAICc relative to null model = -5.56). In the association time model, scaled time with other females was positively associated with offspring survival (OR = 2.27, 95% CI = 0.83 – 6.58), and in the grooming rate model, scaled grooming rate with other females was also positively associated with offspring survival to age 1 (OR = 1.81, CI = 1.11 – 3.28). However, because model fit for the CSI-F model was considerably better, the composite sociality index appears to better capture social dynamics associated with offspring mortality in the first five years of life than do its component parts.

To explore the complex relationships between our predictor terms, CSI-F, and offspring survival to age 5, we fit a path model using our survival to age 5 dataset. As before, this data set included 97 offspring born to 29 mothers between 1980 and 2017. We conducted the analysis using the lavaan package in R^7^, using the default diagonally-weighted least squares estimator.

We specified kin presence (i.e. the presence of a female’s mother, daughter, or maternal sister over age 11), female age, and squared female age as predictor (i.e. exogenous) variables, with female Elo score as an intermediate outcome (i.e. endogenous) variable predicted by kin presence and female age, CSI-F as a second intermediate (i.e. endogenous) variable predicted by kin presence, female age, squared female age, and female Elo score, and offspring survival as a binary outcome (i.e. endogenous) variable predicted by all five other terms (Figure S4). All tests indicated satisfactory model fit (Model 𝜒^2^ = 0.149, Df = 1, *p* = 0.699; RMSEA = 0.000, *p* = 0.732; CFI = 1.00; SRMR = 0.000). We then used the lavaanPlot package in R^8^ to visualize the associated path diagram (Figure S4).

Path analysis revealed that females had higher CSI-F values if they had close female maternal kin in the community, and that young and old females had stronger CSI-F values. CSI-F in the year preceding birth was directly associated with offspring survival to age 5 (Table S3, Figure S4). Furthermore, unlike the survival to age 1 analysis, kin presence was also directly associated with offspring survival to age 5; however, this effect was negative (after accounting for the effect of CSI-F), such that females with female kin in the community had *lower* offspring survival to age 5. However, these females tended to have higher CSI-F values, which was positively associated with offspring survival to age 5.

##### Distribution of dyadic bond strengths

We calculated the strength of each sample female’s dyadic bonds (Dyadic Sociality Indices, or DSIs^9^) using 1) time together and 2) time spent grooming, standardized by their respective mean values in each window. We then averaged these two values together to calculate DSI values for each dyad. This differed from our CSI calculations in the main analysis because a number of female dyads were not observed together in a given one-year window when one or the other was the focal subject, meaning their grooming rates were undefined.

Exploratory analysis revealed that the strongest dyadic bonds tended to be formed with close female kin (Figures S1, S2). We therefore ran a linear mixed model with each female’s top DSI measure in the year before giving birth as the outcome variable, a binary indicator term for kin status (close kin yes/no), maternal Elo score on the date of birth, maternal age, and squared maternal age as predictor terms, and maternal identity as a random intercept term. We found that, among top bonds, bond strength with close female kin was significantly higher than bond strength with other females (𝛽 = 14.35, 95% CI = [10.10 – 18.60]). In contrast, other terms were not associated with top bond strength (maternal Elo score: 𝛽 = 0.55, CI = [-16.79 – 17.90]; maternal age: 𝛽 = -0.40, CI = [-4.38 – 3.57]; squared maternal age: 𝛽 = 1.29, CI = [-0.31 – 2.89]).

##### Comparison of social integration vs. social bond strength as predictors of offspring survival

Different measures of sociality have been found to predict fitness-related outcomes across different studies, with no obvious way to predict a priori which measure is best^10,11^. Therefore, we ran an additional set of analyses comparing our measure of social integration with other females (CSI-F) with measures of the strength of females’ top dyadic social bonds with other females as predictors of offspring survival. We used DSI values based on time together and time spent grooming (described in the previous section) to determine the strength of each females’ top social bond; first, following early work on social bonds and adaptive outcomes^12–14^ we calculated the summed strength of each female’s top 3 bonds (DSI_time_3) in the year preceding giving birth. Second, because the selection of a set number of top bonds for each individual is somewhat arbitrary^15^, and the number of strong or weak bonds females form can be dependent on group size^16^, we also included a measure of the strength of each female’s top bond (DSI_time_1)^10^.

We next calculated DSI values based on time together and grooming *rate*, because 1) this dyadic measure more closely resembled the CSI-F measure and 2) grooming rate was calculable for all dyadic bonds with top bond partners (but not all dyadic bonds, see previous section). From these measures we also determined the summed strength of females’ top 3 bonds (DSI_rate_3) as well as the strength of their top bond (DSI_rate_1) in the year preceding giving birth.

We then re-ran our best-supported model from the main analysis (see main text), replacing the CSI-F measure with each of the four DSI-based measures of social bond strength. We compared these four models to the CSI-F model using AICc as the selection criterion (Table S4). To ensure effect sizes were comparable, we scaled each sociality term by Z-transformation before running each model. The model including CSI-F, our measure of social integration with other females (presented in the main text), remained the best-supported model (model weight = 0.54), while the model including the strength of each female’s top rate-based bond (DSI_rate_1) had similar support (ΔAICc relative to the best-supported model = 1.88; model weight = 0.21). The association between offspring survival and 1) CSI-F and 2) DSI_rate_1 was also similar, although again the former measure was slightly more strongly associated with offspring survival (Table S4).

This suggests that the strength of females’ top social bond was an important component of their overall social integration, and that these two measures are capturing similar social dynamics. As evident in Figure S1, females appear to form one, or sometimes two, especially strong social bonds, producing a highly skewed distribution of bond strengths. In turn, this suggests that it may be difficult to disentangle the effects of females’ strongest bond with those of social integration as distinct components of female sociality, at least in the Gombe population. These results echo those from a recent analysis of data from five species of primates, finding that measures of social bond strength and social integration tend to be highly correlated and likely capture the same biological phenomenon^17^.

##### Comparison with previous studies of offspring survival in the same population

The main and supplementary analyses found no relationship between maternal rank score and offspring survival to age 1 (see main text), and only an indirect relationship between maternal rank score and offspring survival to age 5, via the relationship between maternal rank score and maternal social integration (Table S3; Figure S4). This differs from previous analyses in Gombe, which found that rank predicted offspring survival to age 7^18^ and to age 8^19^. To determine whether different survival ages are the source of the discrepancy between previous and current analyses of Gombe data, we conducted an additional analysis of offspring survival to age 7. We followed the protocol outlined in the methods section of the main text and the Survival to age 5 analysis above. Although offspring whose mothers died between age 5 and age 7 may survive to adulthood, maternal loss remains costly for juveniles in that age range^20^, so we restricted our data set to offspring born to females that survived at least 7 years after giving birth. This resulted in a data set of 84 offspring born to 25 mothers from 1981 to 2016. 38 of the 84 offspring in this sample died before age 7.

As above, we used logistic regressions to model the odds ratios of offspring survival to age 7^1^, using the lme4 and stats packages in R^2,3^. As with the primary analysis, we first constructed a global model containing all non-sociality predictor terms (offspring sex, firstborn status, maternal rank score, maternal age and squared maternal age, maternal immigration status, maternal SIV status, and counts of males and females in the community). As before, the random intercept term for maternal identity did not account for any variance, so we excluded it from future models.

We again used a model comparison procedure, with AICc as the selection criterion, to determine the best set of non-sociality terms for predicting offspring survival to age 7. We used the dredge function in the R package MuMIn^4^ to compare all subsets of the global model, excluding only models from the candidate set that included the squared maternal age term but *not* the corresponding first order term.

We again used the nesting rule to exclude models from the top model set that differed from a better-supported model by the addition of a single term^5,6^, and considered models in the candidate set of best models if they differed from the best-supported model by less than 6 AICc points^5^. The best fitting non-sociality model included three terms: offspring sex, maternal immigration status, and count of sexually-mature females in the community at birth. Notably, maternal rank score did not appear in any of the models in the top model set. Current results therefore differ from earlier analyses of maternal rank score and offspring survival in Gombe even when using the same survival age.

To further investigate the discrepancy in results, we generated a new dataset of 25 offspring born between 1970 and 1979, the period for which we lack grooming data (and we therefore excluded from the main analysis) but for which we can still determine maternal rank score at birth. 10 of these 25 offspring died before age 7. We then compared maternal rank scores for offspring that survived to age 7 with those for offspring that died before age 7 in the two periods. In the earlier period, maternal rank scores differed considerably by offspring survival, with mothers of offspring that survived tending to have higher rank scores than mothers of those that died. However, in the period of the current study, maternal rank score did not differ by offspring survival (Figure S5). Previous analyses of offspring survival in Gombe included data from this earlier period^18,19^, which likely accounts for the differing results. Follow-up work is required to better understand the cause of the changing association between maternal rank score and offspring survival over the *longue durée* of the Gombe project.

#### Supplementary tables

**Table S1:** Maternal kin present in community by maternal residence status

| **Maternal residence status** | **Total mothers** | **Total offspring** | **Total with >= 1 female kin in community** | **Mom has her mom in community** | **Mom has maternal sister in community** | **Mom has adult daughter in community** |
| --- | --- | --- | --- | --- | --- | --- |
| Immigrant | 23 | 51 | 8 | 0 | 2 | 6 |
| Natal | 12 | 53 | 31 | 24 | 12 | 5 |
| Unknown | 2 | 6 | 1 | 0 | 0 | 1 |

Note that some offspring are born to mothers with more than one class of maternal kin present, so row-wise sums of columns 5 to 7 may not match values in column 4.

**Table S2:** List of models and model fit parameters using measures of social integration to predict offspring survival age 5

| **Model** | **df** | **AICc** | **ΔAICc** | **weight** |
| --- | --- | --- | --- | --- |
| Null + CSI-F | 3 | 125.782 | -5.564 | 0.53 |
| Null + CSI-F + CSI-M | 4 | 126.524 | -4.823 | 0.366 |
| Null + gregariousness | 3 | 131.004 | -0.342 | 0.039 |
| Null | 2 | 131.346 | 0 | 0.033 |
| Null + CSI-M | 3 | 131.387 | 0.041 | 0.032 |

Models include all terms from the null model, plus the term(s) described in the model column. **ΔAICc** shows the difference in corrected AIC score between each model and the null model, with negative values indicating models that fit better than the null model.

**Table S3**: Results from path analysis of sociality and offspring survival to age 5, representing a hypothesized set of causal relationships among the maternal social environment, maternal social integration, maternal characteristics, and offspring survival (visualized in Fig. S4). Estimates represent unstandardized path coefficients, and effects whose 95% confidence intervals exclude 0 are bolded.

| **Term** | **Estimate (CI)** | **p-value** |
| --- | --- | --- |
| CSI-F ~ |  |  |
| **Maternal female kin** | **0.44 (0.24 – 0.64)** | **< 0.001** |
| **Maternal age** | **-0.25 (-0.42 – -0.08)** | **< 0.01** |
| **Maternal age^2^** | **0.12 (0.01 – 0.23)** | **0.04** |
| **Maternal Elo** | **0.19 (0.03 – 0.34)** | **0.02** |
| Maternal Elo ~ |  |  |
| Maternal female kin | 0.26 (-0.03 – 0.54) | 0.08 |
| **Maternal age** | **0.80 (0.65 – 0.95)** | **< 0.001** |
| Survival to 5 ~ |  |  |
| **CSI-F** | **0.89 (0.34 – 1.43)** | **< 0.01** |
| **Maternal female kin** | **-0.58 (-1.15 – -0.02)** | **0.04** |
| Maternal age | -0.13 (-0.61 – 0.35) | 0.84 |
| Maternal age^2^ | -0.11 (-0.37 – 0.14) | 0.37 |
| Maternal Elo | 0.31 (-0.13 – 0.75) | 0.17 |

Bold rows indicate significant relationships.

**Table S4:** Comparison of models of social integration and four measures of social bond strength predicting offspring survival to age 1. Each model includes the two terms from the Null (non-social) model (offspring sex and count of sexually mature females present in the community at offspring birth) plus the measure of female sociality listed in the first column. **Sociality term (scaled) OR:** odds ratio indicating the effect of a one-standard-deviation change in the sociality term in each model on odds of offspring survival. **Sociality term (scaled) CI:** 95% confidence interval of the scaled OR.

| **Model** | **AICc** | **ΔAICc** | **weight** | **Sociality term (scaled) OR** | **Sociality term (scaled) CI** |
| --- | --- | --- | --- | --- | --- |
| Null + CSI-F | 112.64 | 0 | 0.54 | 2.03 | 1.15 – 4.07 |
| Null + DSI_rate_1 | 114.52 | 1.88 | 0.21 | 1.85 | 1.03 – 3.84 |
| Null + DSI_rate_3 | 116.26 | 3.62 | 0.09 | 1.54 | 0.92 – 2.90 |
| Null + DSI_time_1 | 116.27 | 3.63 | 0.09 | 1.54 | 0.92 – 3.23 |
| Null + DSI_time_3 | 116.71 | 4.08 | 0.07 | 1.45 | 0.89 – 2.79 |

#### Supplementary figures

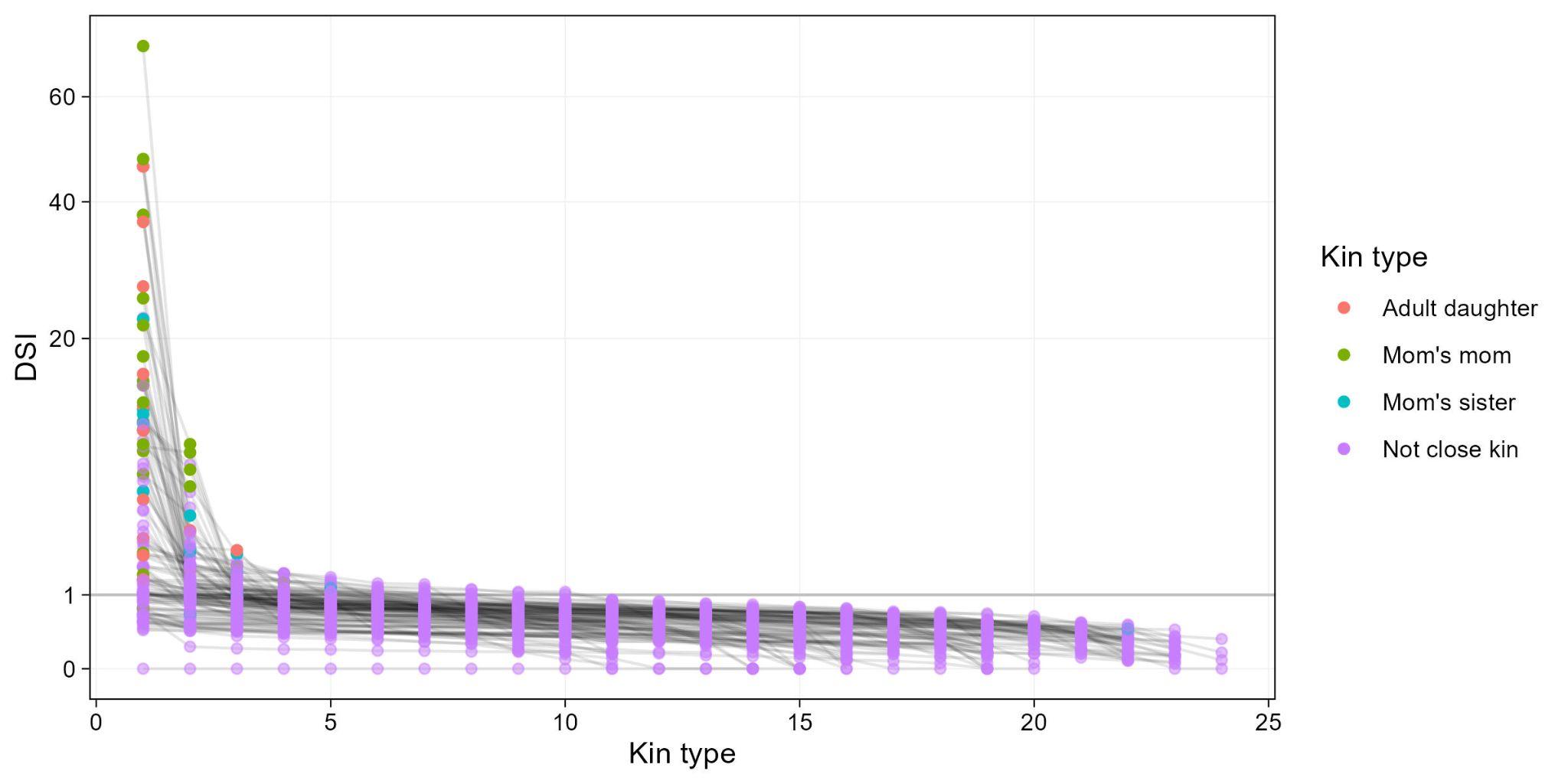

**Figure S1**: Distribution of each target female’s relative dyadic bond strengths (DSIs) in the year preceding giving birth, organized from strongest to weakest bond for each female. Point colors indicate kin class, and faint lines connect values for a given female. A DSI value of 1 represents the mean bond strength among females in a given window. Note the y-axis is on a log scale. Kin types: **Adult daughter**: a target female’s daughter older than age 11 when target female gives birth; **Mom’s mom**: a target female’s mother; **Mom’s sister**: a target female’s maternal sister; **Not close kin**: a female that fits none of these categories.

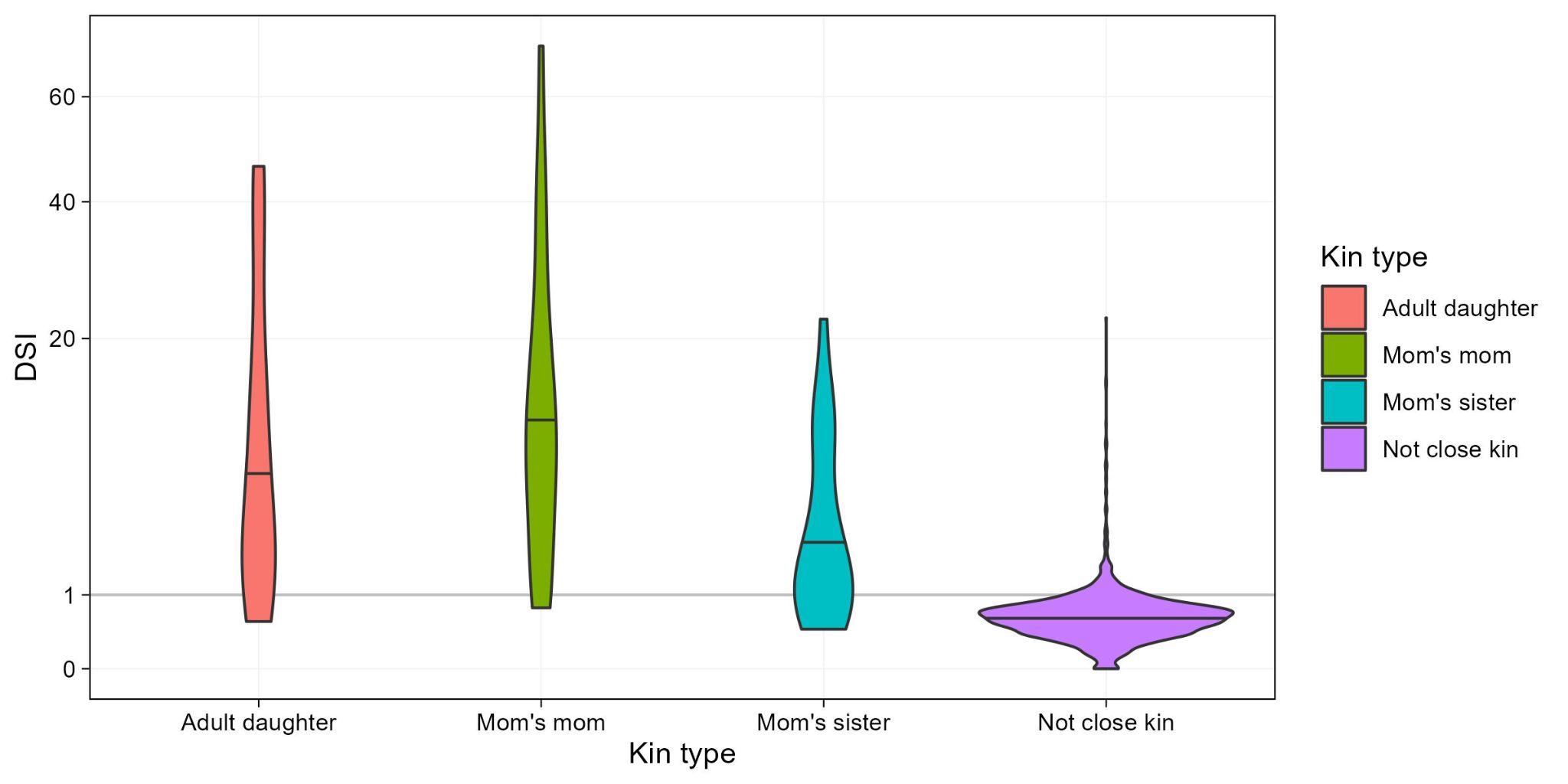

**Figure S2**: Distribution of target female’s relative dyadic bond strengths (DSIs) in the year preceding giving birth, by kin class. Horizontal line in violins indicates the median DSI value for each kin class. A DSI value of 1 represents the mean bond strength among females in a given window. Note the y-axis is on a log scale. Kin types: **Adult daughter**: a target female’s daughter older than age 11 when target female gives birth; **Mom’s mom**: a target female’s mother; **Mom’s sister**: a target female’s maternal sister; **Not close kin**: a female that fits none of these categories.

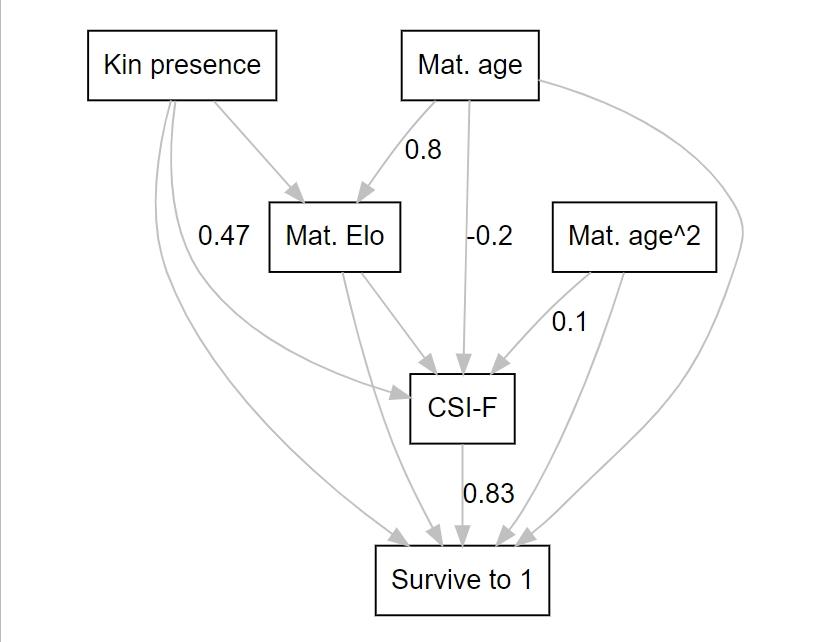

**Figure S3:** Results of path analysis finding age and kin presence effects on CSI-F, and a CSI-F effect on offspring **survival to age 1**, but no direct effects between age, rank, or kin presence terms and offspring survival. Numbers show standardized coefficients for significant effects only (full model results shown in Table 2 in main text). “Mat. age” = maternal age, “Mat. age^2” = squared maternal age, “Mat. Elo” = maternal Elo score at birth, “CSI-F” = maternal Composite Sociality Index with other females.

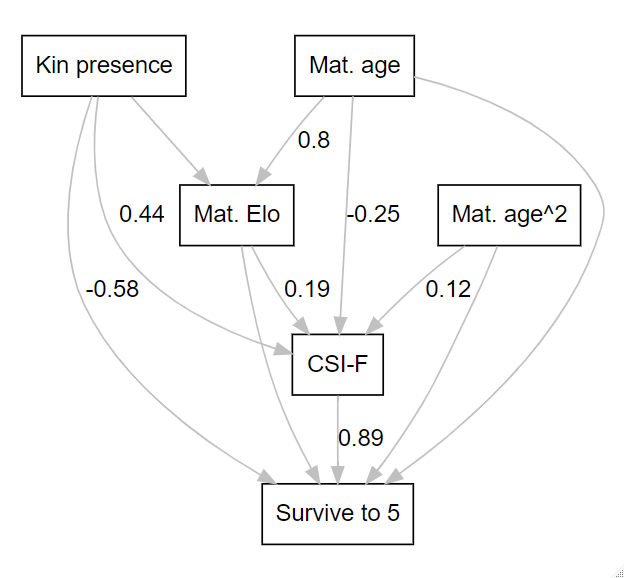

**Figure S4:** Results of path analysis finding age and kin presence effects on CSI-F, and a CSI-F effect on offspring **survival to age 5**, but no direct effects between age, rank, or kin presence terms and offspring survival to age 5. Numbers show unstandardized coefficients for significant effects only (full model results shown in Table S3).

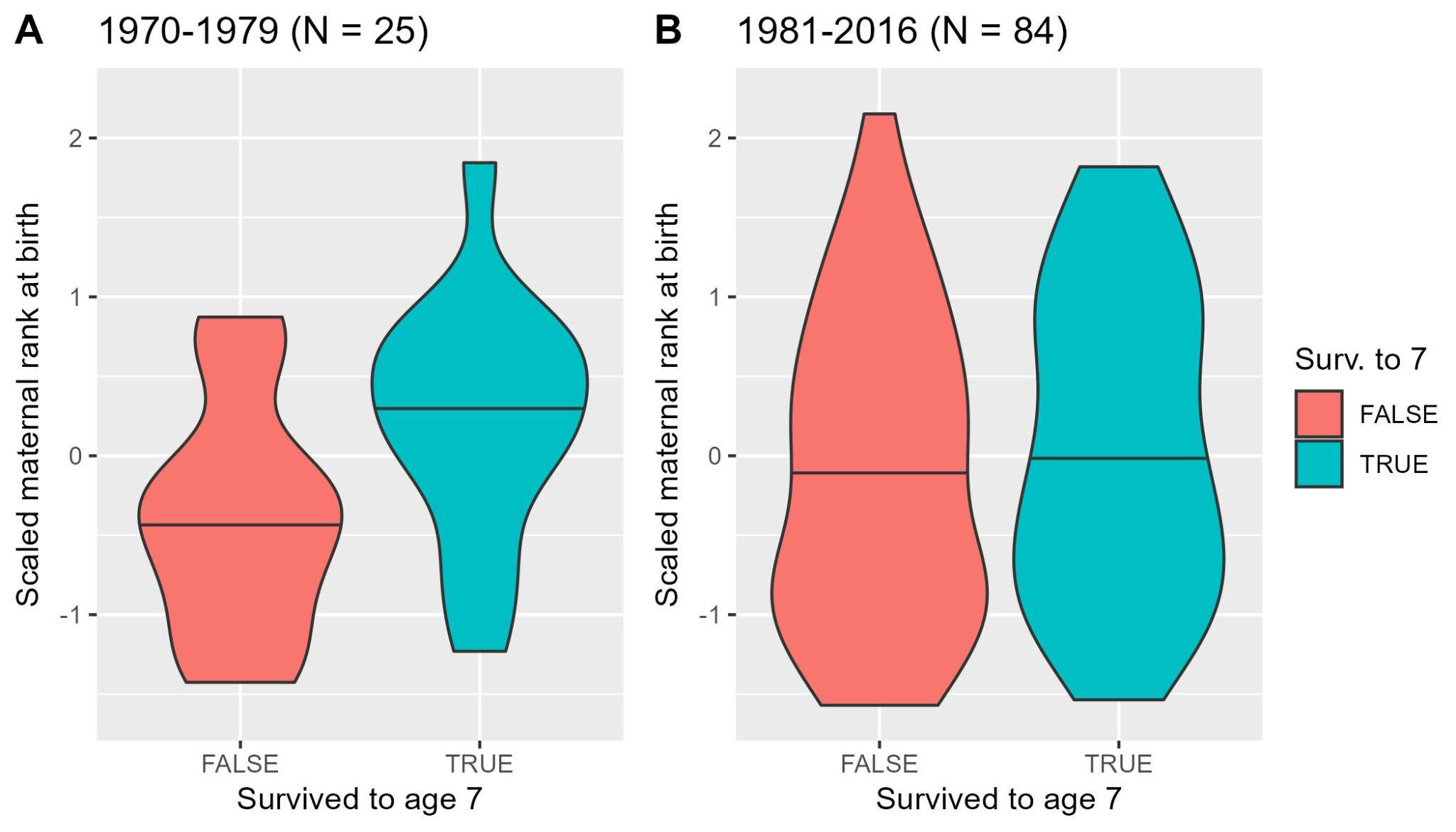

**Figure S5:** Scaled maternal rank at birth vs. offspring survival to age 7 in the 1970s (**A**) and after 1981 (**B**). Horizontal lines in violins indicate the median value of the distribution.
